# “Disruption of Golgi markers by two RILP-directed shRNAs in neurons: a new role for RILP or a neuron-specific off-target phenotype?”

**DOI:** 10.1101/2023.03.08.531742

**Authors:** Chan Choo Yap, Laura Digilio, Lloyd McMahon, Bettina Winckler

**Affiliations:** Department of Cell Biology, University of Virginia, 1340 Jefferson Park Avenue, Pinn Hall 3226, Charlottesville, VA 22908, USA

**Keywords:** gene silencing, neuron, Golgi, endosome, RAB34, RILP, shRNA

## Abstract

In neurons, degradation of dendritic cargos requires RAB7 and dynein-mediated retrograde transport to somatic lysosomes. In order to test if the dynein adaptor RILP (RAB-interacting lysosomal protein) mediated the recruitment of dynein to late endosomes for retrograde transport in dendrites, we obtained several knockdown reagents which had been previously validated in non-neuronal cells. We found that striking endosomal phenotypes elicited by one shRILP plasmid were not reproduced by another one. Furthermore, we discovered a profound depletion of Golgi/TGN markers for both shRILP plasmids. This Golgi disruption was only observed in neurons and could not be rescued by re-expression of RILP. This Golgi phenotype was also not found in neurons treated with siRILP or gRILP/Cas9. Lastly, we tested if a different RAB protein that interacts with RILP, namely the Golgi-associated RAB34, might be responsible for the loss of Golgi markers. Expression of a dominant-negative RAB34 did indeed cause changes in Golgi staining in a small subset of neurons but manifested as fragmentation rather than loss of markers. Unlike in non-neuronal cells, interference with RAB34 did not cause dispersal of lysosomes in neurons. Based on multiple lines of experimentation, we conclude that the neuronal Golgi phenotype observed with shRILP is likely off-target in this cell type specifically. Any observed disruptions of endosomal trafficking caused by shRILP in neurons might thus be downstream of Golgi disruption. Different approaches will be needed to test if RILP is required for late endosomal transport in dendrites. Cell type-specific off-target phenotypes therefore likely occur in neurons, making it prudent to re-validate reagents that were previously validated in other cell types.

## Introduction

Short hairpin (sh) plasmids are powerful tools to study loss-of-function phenotypes by downregulating the expression of proteins of interest and are thus widely used (1). Compared to generating knockout animals or stable knockout lines, the advantages of using sh-plasmids include the ability to rapidly test many candidates, the capacity of relatively acute downregulation, and the ability to achieve partial, timed downregulation for examining the roles of essential genes. In addition, it has been shown that complete knockout phenotypes are sometimes less severe than knockdown approaches because only knockouts but not knockdowns activate cellular compensation pathways, thus dampening loss of function phenotypes (2). The disadvantages of sh-mediated knockdown approaches include incomplete downregulation and the need to carefully distinguish on-target from potential off-target phenotypes (3). This necessary validation is time-consuming and requires tools not always readily available.

We previously showed that downregulation of RAB7 in cultured rat hippocampal neurons impairs maturation of early endosomes (EEs) to late endosomes (LEs), transport of LEs to the soma, and leads to degradation delays of short-lived cargos (e.g. the neuronal membrane receptor NSG2) in dendrites (4). Similar phenotypes were observed when dynein was inhibited (5). Since RILP has a known role as a dynein adaptor for RAB7 (6,7), we wanted to ask if RILP is necessary for degradative flux in dendrites. We already showed that overexpression of RILP causes increased retrograde transport of LEs in dendrites and clustering of LEs and lysosomes in the soma (5), suggesting the hypothesis that RILP serves as the endogenous dynein adaptor for late endosomes.

Since tools to downregulate RILP levels have already been published, we started with the same reagents, including two different shRILP plasmids (8–10) and a RILP-directed siRNA (11). In the course of using the shRILP reagents in our own hands, we discovered unanticipated loss of Golgi markers. These Golgi phenotypes were not seen in non-neuronal cell lines and were not apparent in neurons with siRILP or gRILP/Cas9. We were also unable to rescue the Golgi phenotypes with re-expressed RILP. Since RILP also interacts with the Golgi-associated RAB34 (12), we asked if the Golgi phenotypes might reflect RAB34-dependent RILP functions. Expression of high levels of dominant-negative RAB34 in neurons led to fragmentation of Golgi but did not phenocopy the shRILP Golgi phenotype. Our data is most consistent with the conclusion that the Golgi phenotype is a neuron-specific off-target effect. This off-target neuronal phenotype complicates conclusions about RILP function in neurons based on these reagents. Since our data show that interference approaches based on some sh-plasmids can manifest off-target phenotypes only in some cell types, validation experiments need to be carried out in the same cell type as the actual experiment.

## Results

### Two published short hairpin plasmids targeting rat RILP give different phenotypes with respect to NSG2 degradation

Inhibition of dynein and expression of RILP-Ct (which no longer binds to dynein complex components DLIC/p150-glued; Fig. 1A) impaired degradation of the short-lived cargo NSG2 in soma and dendrites of cultured rat hippocampal neurons (5). We thus wanted to determine if reducing RILP levels in neurons phenocopied this degradative impairment of NSG2. Downregulation of RILP using siRNA has been published for HeLa cells and causes delayed degradation of EGFR, prolonged accumulation of EGFR in early endosomes (EEs), and accumulation of late endosomal proteins, such as LAMP1 and CD63 (13). shRILP plasmids have also been used in rodent non-neuronal cell lines (Suppl. Table 1) to study the roles of RILP in secretory granules (8). Validation experiments routinely include Western blots or RT-PCR (to confirm on-target efficacy) and using more than one shRILP or siRILP (to guard against off-target effects). In neurons, two shRILP plasmids were previously used to study the role of RILP in axonal transport of signaling endosomes (mouse SCG neurons; (10)) and of autophagosomes (rat cortical neurons; 14). We thus obtained these same shRILP plasmids (designated #1 and #2; Fig. 1B). Both of these sh-plasmids target sequences in the 3’ UTR of the rat or mouse RILP genes. The rat and mouse sequences in the 3’UTR are identical for the sequence targeted by shRILP#1. siRNA against RILP was also used in rat hippocampal neurons to study axonal autophagosome transport (11). Similarly to the shRILP plasmids, the published siRNA is directed at a 3’ UTR sequence (Fig. 1B).

**Figure 1:**
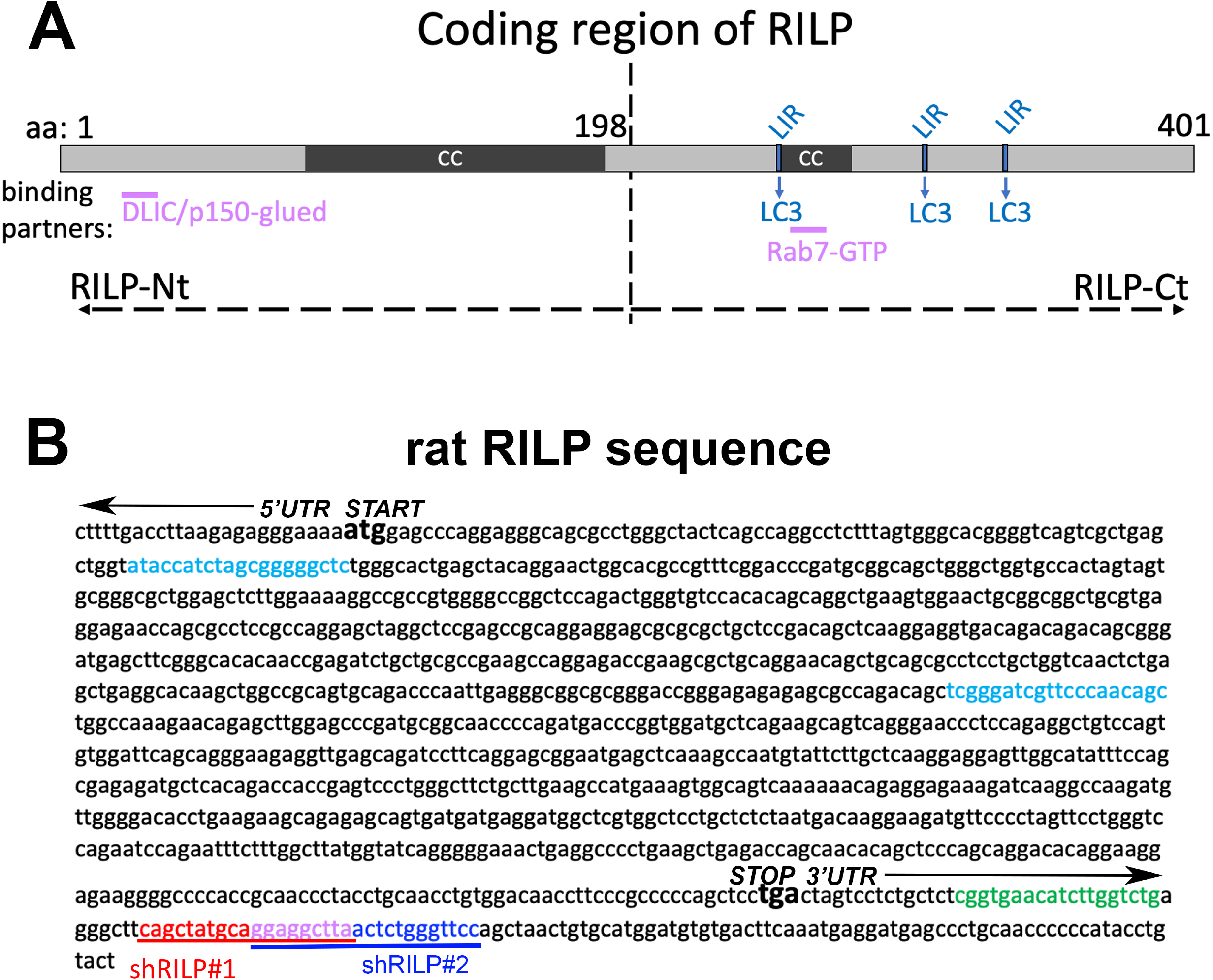
Diagram of RILP protein structure and sequence. (A) Human RILP domain structure is indicated with RILP-Nt encompassing amino acids 1-198 and RILP-Ct encompassing amino acids 199-401. The two coiled coil regions are indicated as CC. The three LC3-binding regions are indicated as LIR. Known binding sites to dynein/dynactin (DLIC and p150) and Rab7 are marked by purple lines. (B) Nucleotide sequence of rat RILP is shown. Start and stop codons as well as 3’ UTR are indicated. Sequences targeted by interference approaches are indicated: shRILP#1 (red) and #2 (dark blue) are partially overlapping in the 3’UTR. siRILP (green) is upstream in the 3’UTR. gRILP/Cas9-targeted sequences are located in the coding sequence and indicated in color (light blue). See Supplemental Table1 for additional details.

Our previous work has shown that the short-lived dendritic cargo NSG2 accumulates 1.5- to two-fold basally when RAB7 or dynein function are inhibited (5). NSG2 levels are greatly reduced (by about 70%) in the soma when new protein synthesis is inhibited for 4 hours using cycloheximide (CHX), but RAB7 or dynein inhibition largely prevent the drop in NSG2 during CHX, implicating RAB7 and dynein in normal degradation of NSG2. In order to test if downregulation of RILP affected NSG2 levels at steady state (T=0) or delayed NSG2 degradation during CHX chase (T=CHX 4h), we transfected cultured rat hippocampal neurons at day 5 in vitro (DIV5) with shControl or shRILP (Fig. 2A-D) and determined the levels of NSG2 six days later (DIV11) either basally (T=0; Fig. 2A,C,E,G) or after 4 hours of CHX (Fig. 2B,D,F,H). Similarly to RAB7 inhibition or RILP-CT overexpression (5), shRILP#1 expression led to accumulation of NSG2 at baseline (T=0; Fig. 2E) and to decreased degradation after 4h CHX (Fig. 2F), suggesting degradation is impaired with reduced RILP levels. When we used a different shRILP plasmid (#2) to test if the observed phenotype was the same, we could not phenocopy the degradation delay seen with shRILP#1 (Fig. 2C,D,G,H).

**Figure 2:**
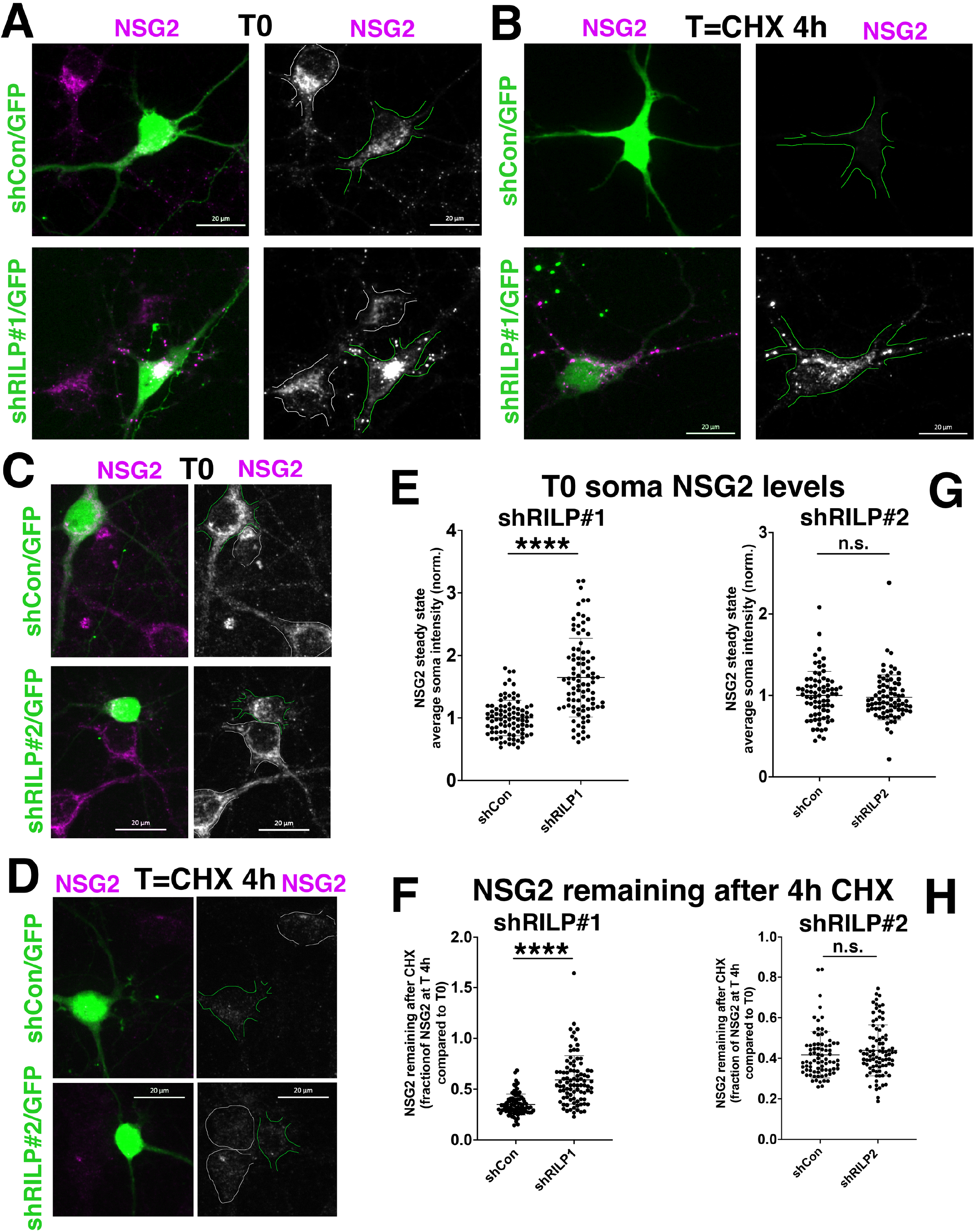
shRILP#1 and #2 give different phenotypes for degradation delay in dendrites. Levels of the short-lived dendritic receptor NSG2 in the soma were determined at base line (T0; A,C,E,G) and after 4 hours of cycloheximide treatment (T=CHX 4h; B,D,F,H) for shRILP#1 vs shCon (A,B,E,F) and shRILP#2 vs shCon (C,D,G,H). shCon is a non-targeting control construct. shRILP#1 but not shRILP#2 led to increased levels of NSG2 at T0 and delayed degradation at T=CHX 4h, thus giving inconsistent results. NSG2 staining alone is shown in the black and white panels. The transfected cell is outlined in green. Untransfected cells in the same field are outlined in white. Rat hippocampal neurons were transfected in culture at DIV5 and fixed at DIV11. Quantification is from 75-95 cells from 3 independent cultures. Mann Whitney statistical test was used.

### Inconsistent phenotypes for endosomal markers in neurons expressing shRILP#1 or shRILP#2

In parallel to NSG2 staining, we also stained against multiple proteins found in early endosomes (EEA1; Fig. 3A,D), late endosomes (RAB7; Fig. 3B,E) and endosomes decorated with retromer (VPS35; Fig. 3C,F) in neurons transfected with shControl, shRILP#1 or −#2. Again, we observed different phenotypes for the two shRILP constructs. shRILP#1 showed strong accumulation of EEA1 (Fig. 3A), and moderate accumulation of RAB7 (Fig. 3B) and of VPS35 (Fig. 3C). In contrast, we did not observe any intensity changes in EEA1 (Fig. 3D), RAB7 (Fig. 3E), or VPS35 (Fig. 3F) for shRILP#2 compared to shControl.

**Figure 3:**
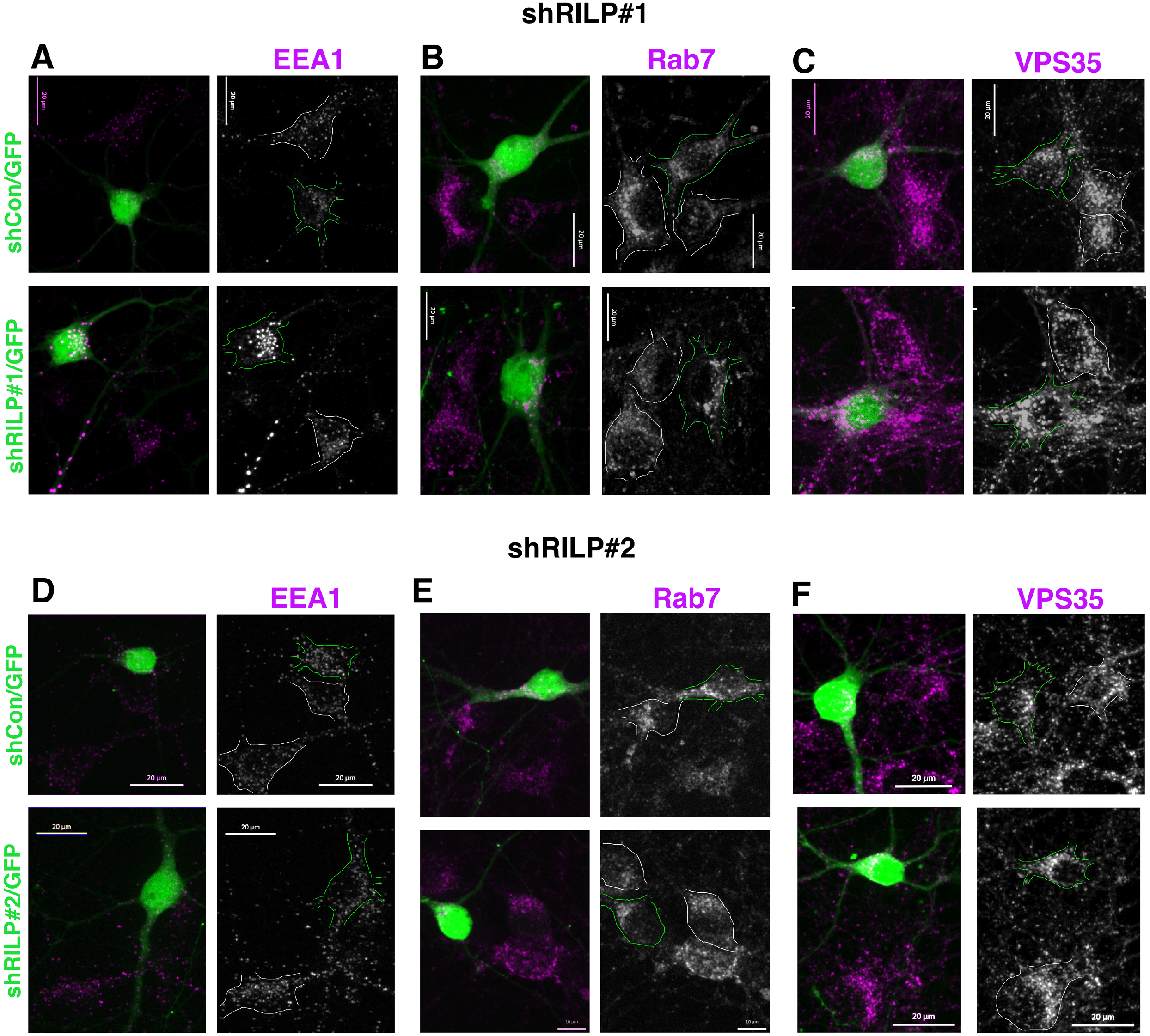
Rab7, EEA1 and Vps35 accumulate in shRILP#1- but not shRILP#2-expressing in neurons. DIV11 at hippocampal neuronal cultures were stained for different endosomal markers after 6 days of expressing shRILP#1 (A-C) or #2 (D-F). Counterstains included EEA1 (A,D), Rab7 (B,E), and Vps35 (C,F). shCon is a non-targeting control construct. Marker staining alone is shown in the black and white panels. The transfected cell is outlined in green. Untransfected cells in the same field are outlined in white.

One possible explanation for the discrepant phenotypes could be better efficacy of shRILP#1 in downregulating endogenous RILP levels, thus giving rise to the different penetrance of the observed phenotypes. shRILP#1 was shown by Western blot to reduce RILP levels by 80% in mouse insuloma cultures (8). shRILP#2 efficacy was validated in previous publications by Western blot and by qRT-PCR in rat C6 glioma cells, leading to 60-70% downregulation (14). In order to test whether shRILP#1 was more potent than shRILP#2 in our rat neuronal cultures, we obtained two commercial antibodies against RILP (ab128616 and ab140188). Our validation of species specificity of these two reagents shows that they only recognize overexpressed tagged versions of human RILP (Suppl. Fig. 1A,B), but not of rat (Suppl. Fig. 1C,D) or mouse RILP (Suppl. Fig. 1E,F) by both immunofluorescence (Suppl. Fig. 1A-F) and Western blot (Suppl. Fig 1G-I) (see also 15). We can therefore not evaluate the relative on-target efficacy of the two shRILP constructs in rat neurons using these antibodies.

### Shared Golgi phenotypes for shRIL#1 and −#2 in neurons, but not non-neuronal cells

Since RILP has additionally been implicated in some Golgi/TGN processes via its interaction with the Golgi-resident RAB34 (12), we also stained against LAMP1 which is trafficked to late endosomes (LEs) and lysosomes from the TGN. LAMP1 staining in the soma was slightly decreased for both shRILP#1 (Fig. 4A,B,E) and shRILP#2 (Fig. 4C-E). This might correspond to an on-target phenotype since it is shared by the two shRILP constructs. We also noticed that soma size was strikingly smaller in shRILP#2 expressing neurons (Fig. 4F) but was not affected by shRILP#1.

**Figure 4:**
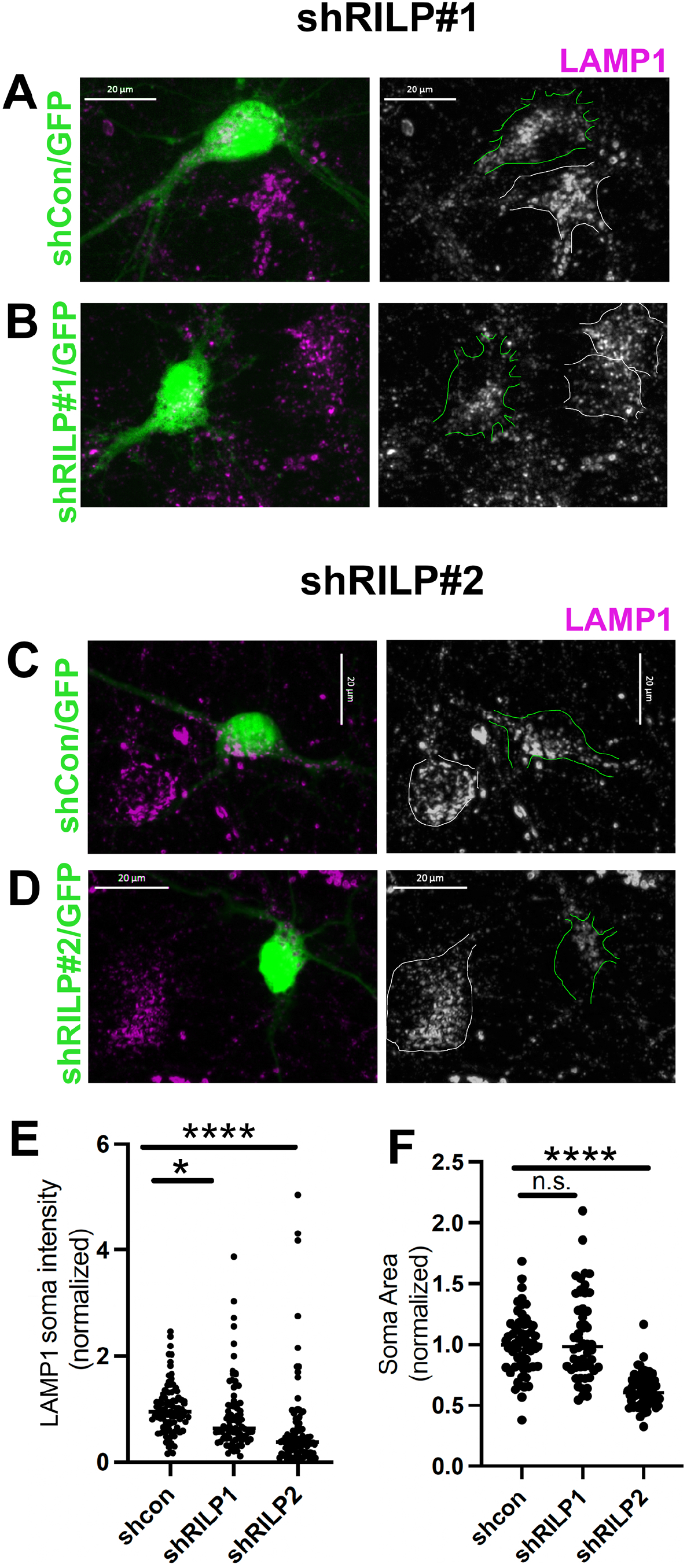
LAMP intensity is decreased in neurons for both shRILP#1 and #2. DIV11 at hippocampal neuronal cultures were stained for LAMP1 after 6 days of expressing shRILP#1 (B) or #2 (D). shCon-GFP is a non-targeting control construct (A,C). LAMP1 staining alone is shown in the black and white panels. The transfected cell is outlined in green. Untransfected cells in the same field are outlined in white. (E) Quantification of LAMP1 staining intensities in the soma of cells transfected with shCon, shRILP#1, or #2. Measurements were normalized to shCon. Quantification is from 81-96 cells from 3 independent cultures. Kruskal Wallis statistical test was used. (F) Quantification of soma size of cells transfected with shCon, shRILP#1, or #2. Quantification is from 54-65 cells from 2 independent cultures. Kruskal Wallis statistical test was used. Measurements were normalized to shCon.

We additionally evaluated Golgi (GM130, a scaffolding protein) and TGN markers (TGN38, a membrane protein shuttling between TGN and endosomes) in rat neuronal cultures transfected with shRILP#1, −#2, or shControl plasmids. We find that both GM130 (Fig.5 A-D) and TGN38 (Fig.5 E-H) staining is greatly reduced (quantification in Fig.5 D,H) compared to shControl or untransfected neurons in the same field. We thus wondered if the shRILP Golgi/TGN phenotype might be on-target since reduced levels of TGN38 and LAMP1 phenotypes are observed for both shRILP plasmids, suggesting a possible role for RILP in trafficking of TGN38 and LAMP1.

**Figure 5:**
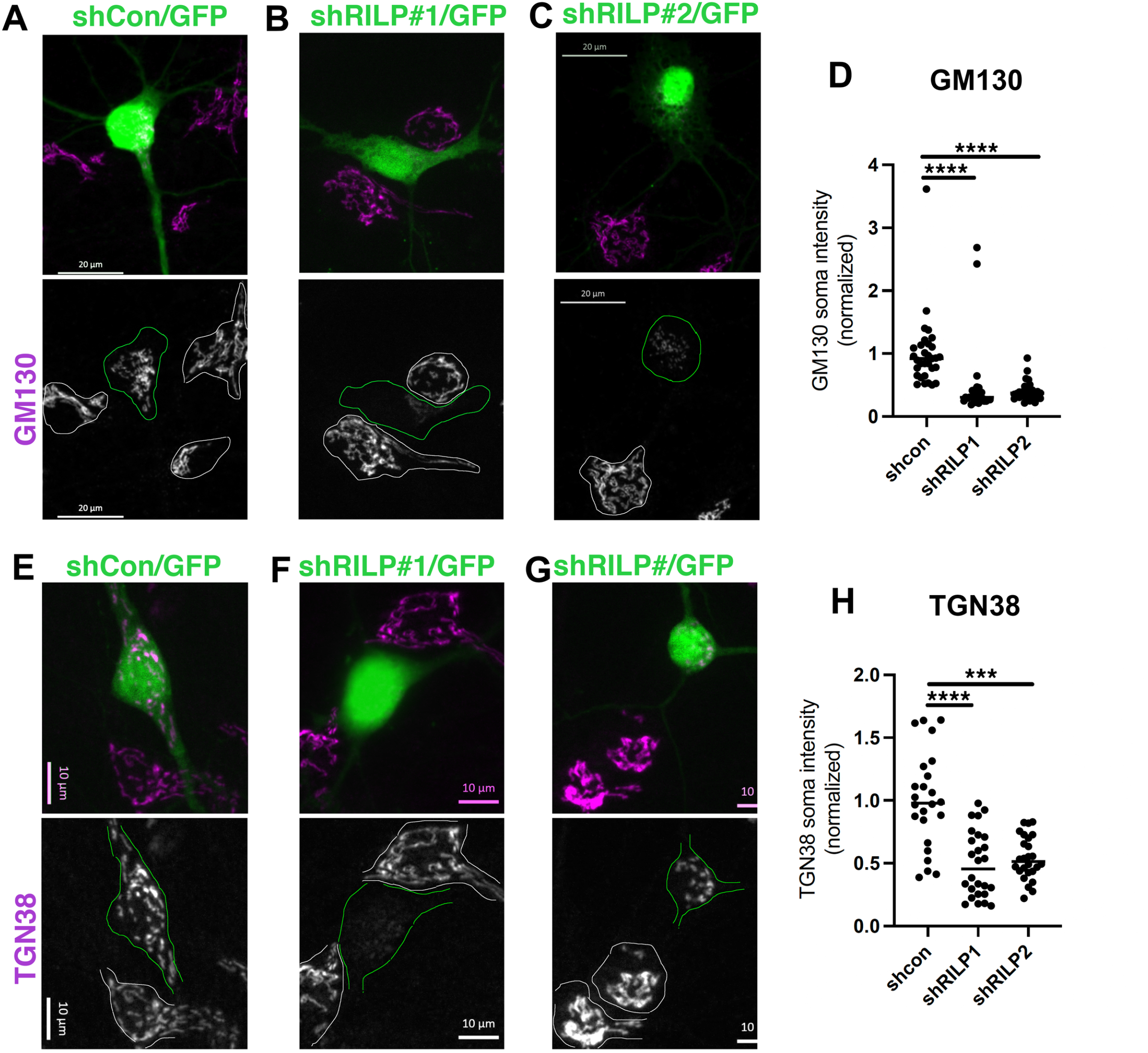
Golgi markers are decreased for both shRILP#1 and #2 in neurons. DIV11 at hippocampal neuronal cultures were stained for GM130 (A-C) of TGN38 (E-G) after 6 days of expressing shCon (A,E), shRILP#1 (B,F) or #2 (C,G). Marker staining alone is shown in the black and white panels. The transfected cell is outlined in green. Untransfected cells in the same field are outlined in white. (D,H) Quantification of GM130 (D) and TGN38 (H) staining intensities in the soma of cells transfected with shCon, shRILP#1, or #2. Quantification shown is from 24-31 cells of one experiment. The experiment was repeated 2-3 times. Kruskal Wallis statistical test was used. Measurements were normalized to shCon.

RILP is widely expressed and has been implicated in trafficking to lysosomes in several non-neuronal cell types (7, 13, 16). A Golgi phenotype for downregulating RILP was not previously reported. We thus determined if Golgi/TGN markers were also reduced in a nonneuronal rat cell line, NRK (normal rat kidney) cells. Surprisingly, GM130 staining appeared entirely normal in NRK cells transfected with shRILP#1 plasmid (Fig. 6A), comparable to nontransfected cells in the same field or cells transfected with shControl plasmid. To assess the endosomal phenotypes we observed with shRILP#1 in neurons (Fig. 3), we also stained transfected NRK cells against RAB7 (Fig. 6B), VPS35 (Fig. 6C), and LAMP1 (Fig. 6D). No changes were observed for any of the markers in NRK cells expressing shRILP#1 compared to nontransfected cells or cells expressing shControl. Unexpectedly then, two different shRILP plasmids cause Golgi phenotypes only in neurons, but not in NRK cells. We then tested if we could rescue GM130 staining levels in neurons by simultaneous co-transfection of shRILP#1 or #2 with myc-HumanRILP in neurons (Fig. 7). GM130 levels were still greatly reduced even when myc-HumanRILP was re-expressed (Fig. 7 D,F). We could thus not convincingly demonstrate that the Golgi phenotype is on-target in neurons.

**Figure 6:**
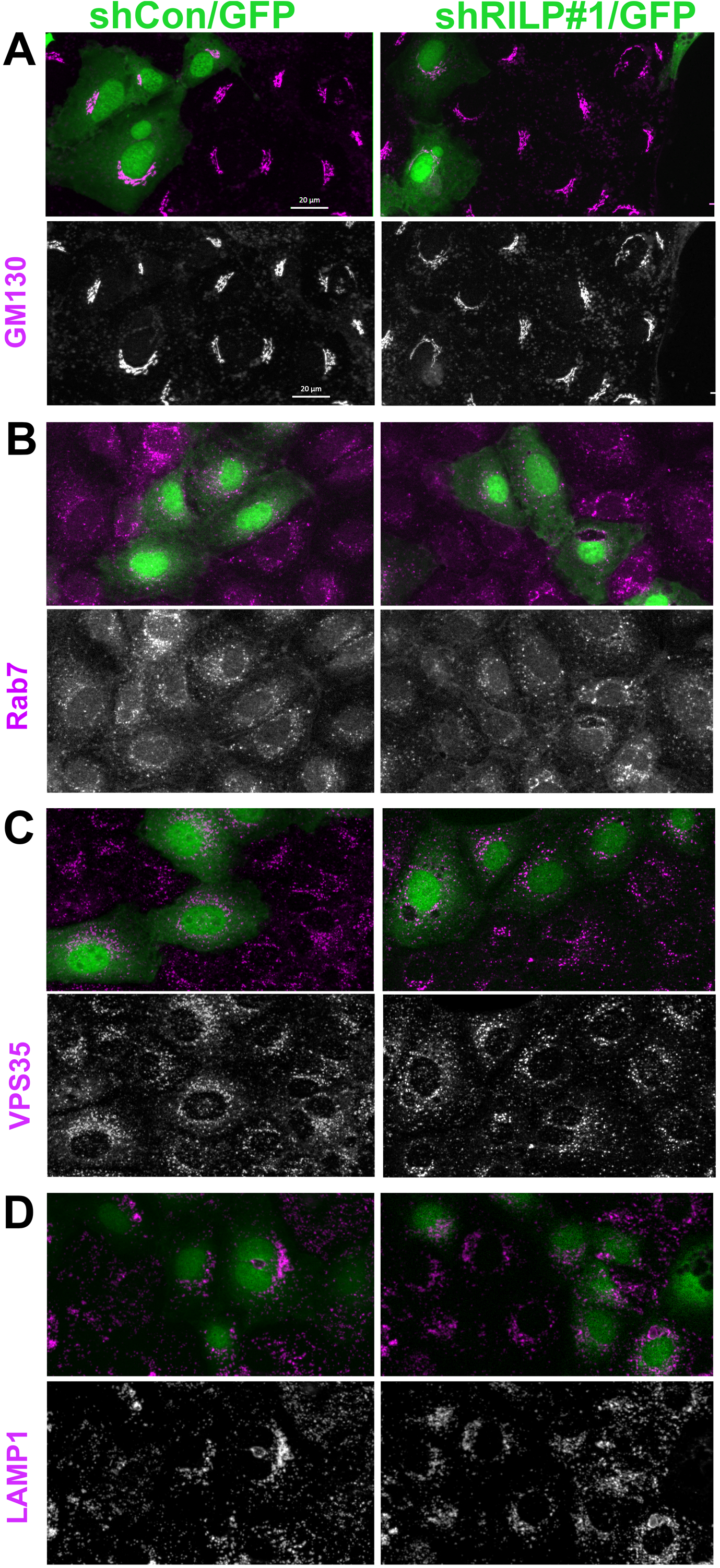
Normal rat kidney (NRK) cells transfected with shRILP#1 do not show Golgi or endosome marker changes. NRK cells were stained for different organelle markers after 5 days of expressing shCon (left panels) or shRILP#1 (right panels). Counterstains included GM130 (A), Rab7 (B), Vps35 (C), and LAMP1 (D). shCon-GFP is a non-targeting control construct. Transfected cells (green) do not show changed marker distribution or intensities (purple). Marker staining alone is shown in the black and white panels.

**Figure 7:**
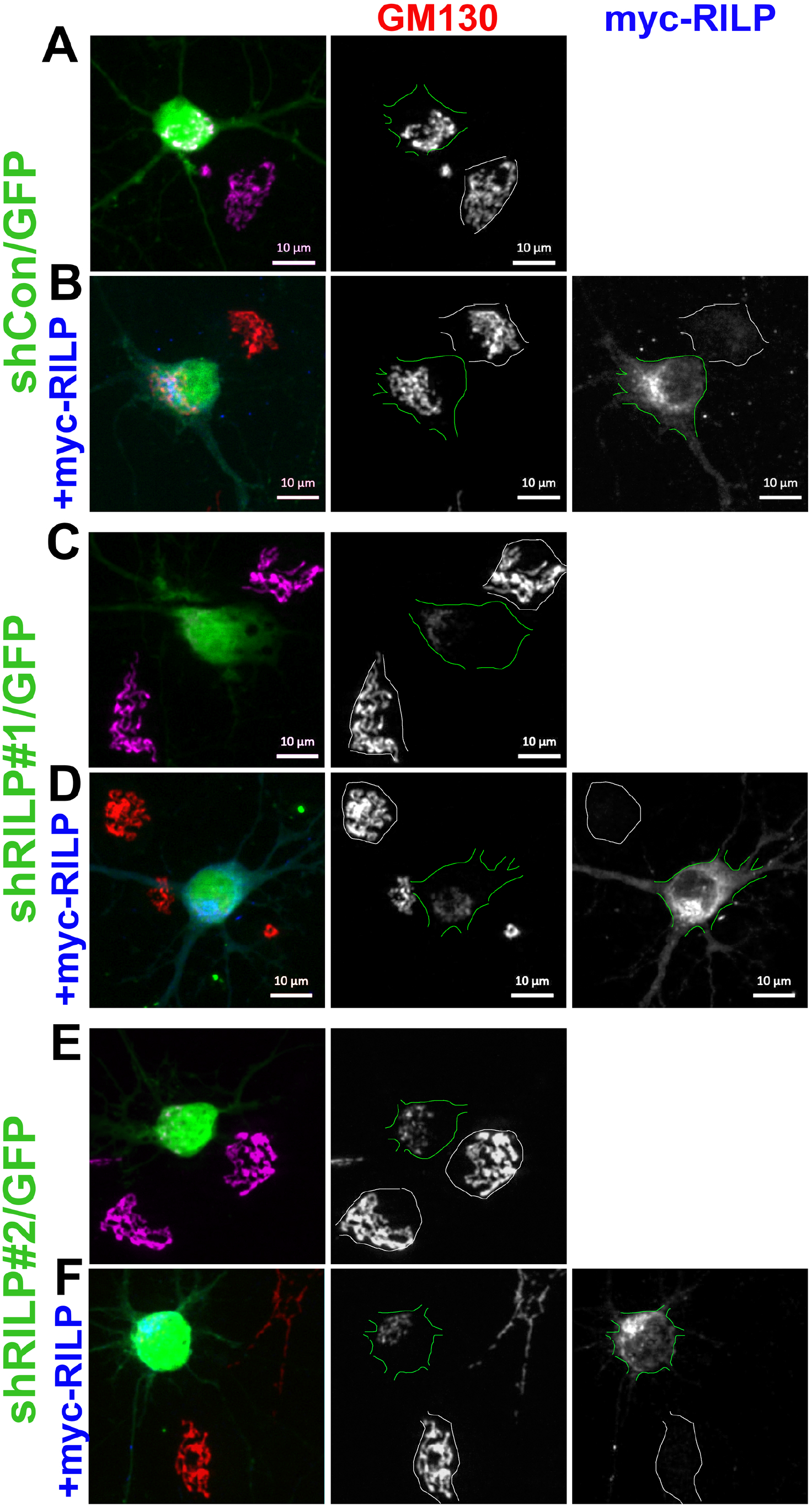
Loss of GM130 from shRILP#1 or #2 expression cannot be rescued by WT RILP re-expression. DIV11 at hippocampal neuronal cultures were stained for GM130 (red) after 6 days of expressing shCon (A,B), shRILP#1 (C,D) or #2 (E,F). In B,D,F, myc-HmRILP was co-transfected (blue) to assess ability to rescue loss of GM130 staining. Marker staining alone is shown in the black and white panels. The transfected cell is outlined in green. Untransfected cells in the same field are outlined in white. Re-expression of myc-HmRILP did not rescue loss of GM130 caused by shRILP#1 and #2.

Lastly, we obtained siRILP (Fig. 1B; Fig. 8A,B) and guide RNA targeting coding sequence of RILP (gRILP) (Fig. 1B; Fig. 8C-F). Cas9/gRILP transfection potently inhibited the expression of Rat RILP-myc from a co-transfected plasmid (Fig. 8C,D). Since the siRNA is directed against 3’UTR, we could not confirm its efficacy using plasmid-driven RILP expression. Published data report a 25% downregulation of RILP after 36 hours in rat PC12 cells by Western blot (11). We observed no Golgi disruption phenotypes in transfected neurons in either experimental paradigm (Fig. 8B,F).

**Figure 8:**
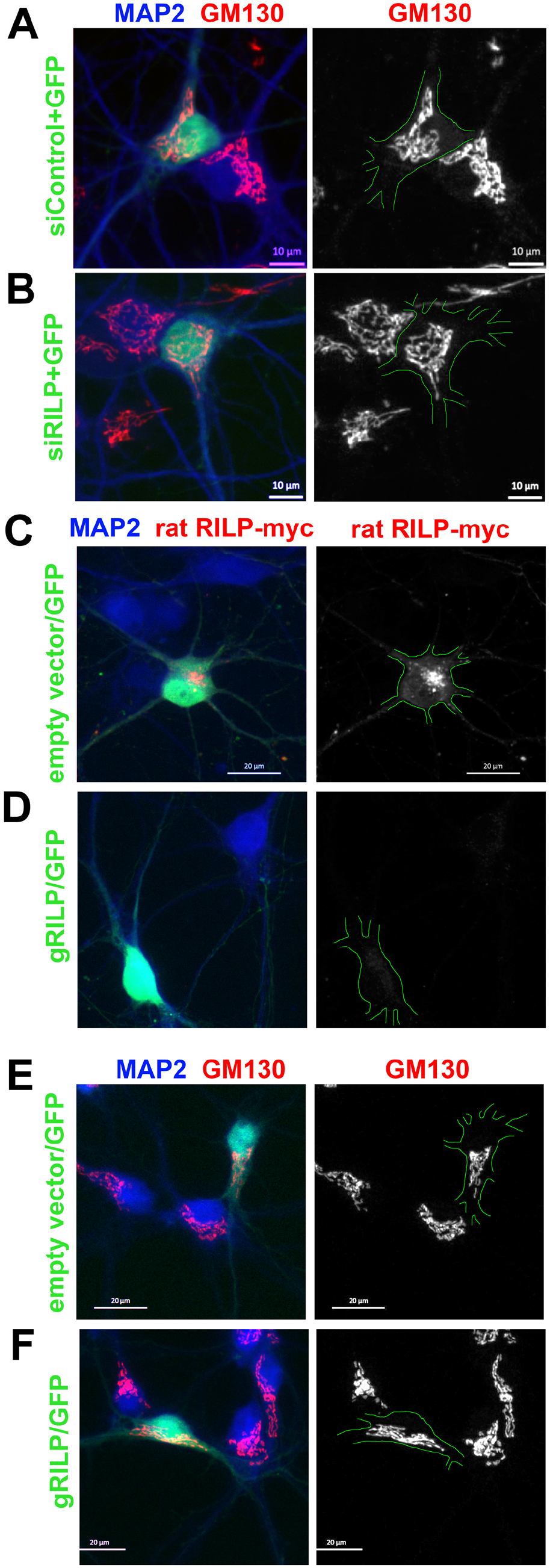
No disruption of Golgi-associated GM130 in RILP downregulation using siRNA or CRISPR/Cas9. (A,B) DIV11 at hippocampal neuronal cultures were stained for GM130 (red) after 6 days of expressing siCon + GFP (A) or siRILP + GFP (B). MAP2 (blue) identifies dendrites. Transfected cells express GFP (green). Transfected cells are outlined in green in the black and white panels showing the GM130 channel. (C,D) DIV11 hippocampal neuronal cultures transfected with rat RILP-myc and either empty vector pDG458 (C) or gRILP (D) were stained against MAP2 (blue) to identify dendrites and anti-myc antibodies (red) to detect RILP-myc. No expression of RILP-myc is seen in cells expressing gRILP, indicating effective targeting of RILP sequence. Transfected cells express GFP and are outlined in green in the black and white panels showing the RILP-myc channel. (E,F) DIV11 hippocampal neuronal cultures transfected with empty vector pDG458 (E) or gRILP (F) for 6 days were stained against MAP2 (blue) to identify dendrites and anti-GM130 antibodies (red). GM130 staining was not affected by gRILP expression. Transfected cells express GFP and are outlined in green in the black and white panels showing the GM130 channel.

### High expression of a dominant-negative Rab34 protein in neurons causes fragmentation of Golgi markers

In order to test in a different way whether RILP downregulation might lead to Golgi phenotypes in neurons as an on-target effect, we decided to test a possible RAB34-RILP-Golgi link. RAB34 also co-immunoprecipitates with RILP (12). RAB34 is a Golgi-localized RAB which is thought to regulate localization of lysosomes to the peri-Golgi region via RILP (12). Expressing of Golgi-localized WT RAB34 but not of dominant-negative (DN) RAB34-T66N in HeLa cells leads to clustering of lysosomes near the Golgi. We transfected rat hippocampal neuronal cultures with GFP, GFP-RAB34, or GFP-RAB34-DN and stained for GM130 (Fig. 9A-C). As reported in HeLa cells, RAB34 staining was relatively enriched on GM130-positive Golgi membranes (Fig. 9B). To better appreciate the relative enrichments of GM130 and GFP-Rab34, we generated a line scan through the soma (Fig. 9B’). Both markers show relative enrichment and depletion in similar patterns with GFP-RAB34 being more diffusely distributed. Overexpression of GFP-RAB34 did not change GM130 distribution. GFP-RAB34-DN, on the other hand, occasionally (~20% of cells) caused apparent fragmentation of GM130-positive Golgi stacks (Fig. 9C), especially in cells expressing very high levels of GFP-RAB34-DN. These cells often started to look unhealthy. Cells expressing lower levels of GFP-RAB34-DN showed no phenotypes and were indistinguishable from untransfected neurons (^~^80% of cells; not shown). Unlike the findings in HeLa cell (12), lysosome clustering (visualized with LAMTOR4 staining) was not observed by WT RAB34 overexpression in neurons (Fig. 9A-C). Interestingly, LAMTOR4 intensity was decreased in neurons expressing high levels of GFP-RAB34-DN (Fig. 9C), suggestive of a role for RAB34 in trafficking of lysosomal components. More work is needed to fully explore the role of RAB34 in membrane trafficking in neurons.

**Figure 9:**
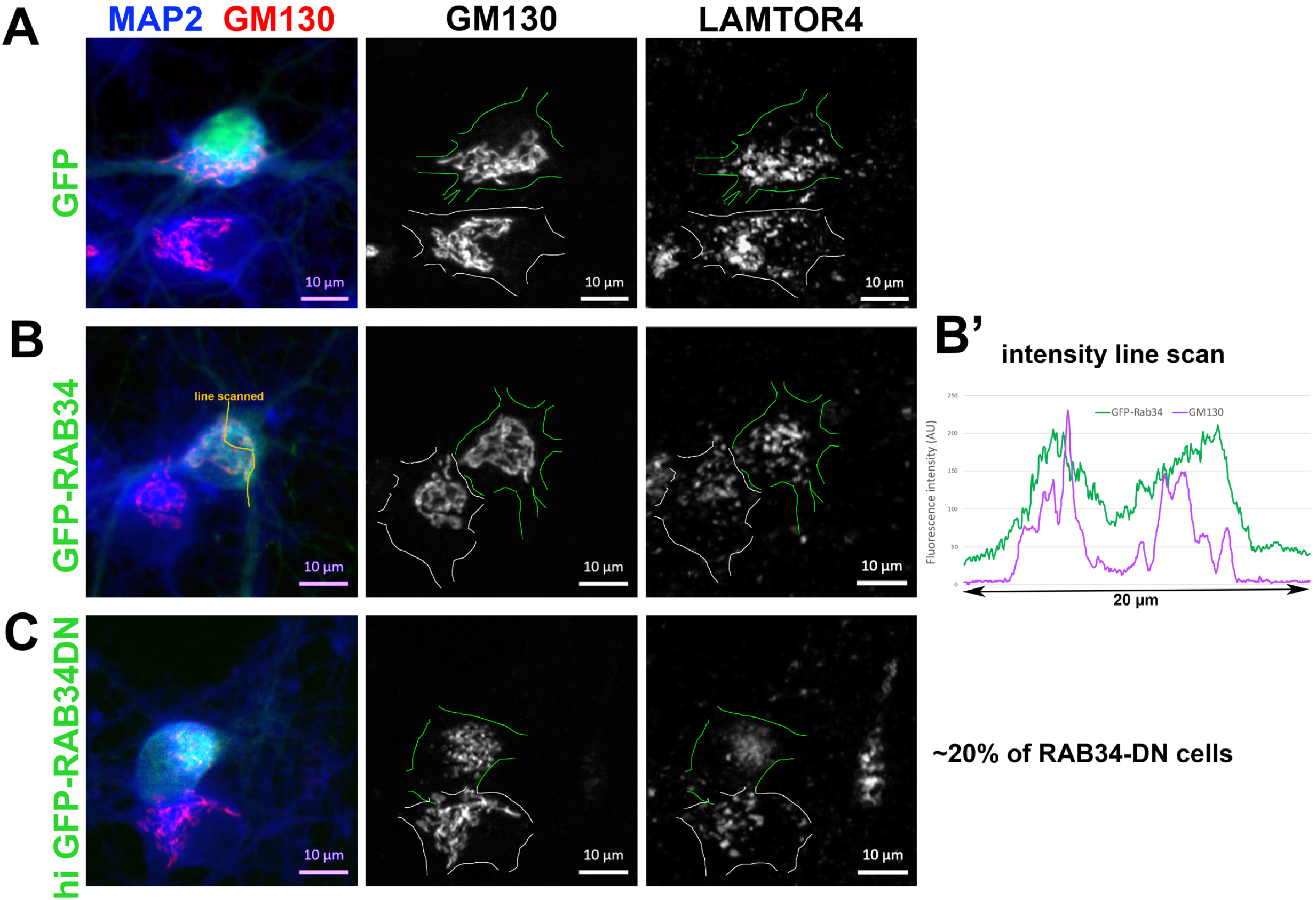
Golgi phenotype caused by Rab34-DN is distinct from the Golgi phenotype caused by shRILP. DIV9 rat hippocampal neuronal cultures were stained for GM130 (red, middle column) or LAMTOR4 (right column) after 48 hours of expressing GFP (A), GFP-Rab34 (B) or high levels of GFP-Rab34-DN (C). GFP-Rab34 localizes to similar cellular regions as GM130 as seen in the linescan in B’. The transfected cell is outlined in green. Untransfected cells in the same field are outlined in white. LAMTOR4 is not clustered by GFP-Rab34 overexpression. High expression of GFP-Rab34-DN in about 20% of cells causes fragmented GM130 staining.

## Discussion

Validation of reagents is critical. Conclusions based on unvalidated reagents might not hold and sow confusion in the field. But how exactly to validate and how extensively is not clear. For instance, is it necessary to rerun validations for reagents for which careful validation has already been published? In this paper, we report a careful validation of several reagents used for studying the function of the RAB7 effector RILP in human cell lines and rodent neurons. For shRILP#1, careful validation was published in rodent insuloma cells (8). These included demonstration of on-target downregulation of endogenous RILP and controls guarding against off-target effects (multiple shRILP constructs giving the same phenotype and phenocopying of the shRILP phenotype by gRILP/Cas9 knockout and two interfering cell penetrating peptides). shRILP#2 was validated previously in rodent glioma cells for on-target efficacy of downregulation using RT-PCR (14). Our work reported here strongly suggests that two short hairpin plasmids directed against the RAB7 effector RILP elicit neuronal-specific off-target effects. Our findings thus argue that knockdown reagents previously validated in other cell types need to be re-evaluated in the experimental cell type of each subsequent study.

Knocking down protein expression with short interfering RNAs (either siRNA or short hairpins) is a powerful approach to evaluate loss-of-function phenotypes for many proteins. Even though knockdown is usually incomplete, phenotypes are often observed giving important insights into protein function. Knockdown is relatively inexpensive and rapid, allowing more acute interference compared to knockout. In addition, in some instances knockout triggers more extensive activation of compensatory pathways, obscuring phenotypes (2, 17). The main caveat against knockdown approaches is the possibility of off-target effects whereby additional mRNAs are also downregulated, microRNAs are targeted, or microRNA processing machinery is saturated (3, 18–21). It is worth pointing out that knockout approaches (especially CRISPR/cas9) are also potentially confounded by their own off-target effects.

The gold standard for validating a knockdown/knockout reagent is to use more than one sequence to be targeted for the same gene and to perform rescue experiments. If two or more knockdown/knockout reagents result in the same phenotype, it is more likely that it is on-target. Since it is basically always possible to test more than one knockdown plasmid, using multiple sh-plasmids is almost always carried out as validation controls. In fact, commercially supplied knockdown reagents usually contain pools three or four distinct siRNAs or sh-plasmids. If possible, the phenotype should also be rescued by re-expression of the targeted protein. Rescue is not always complete, but even a partial rescue lends strength to the argument that the observed phenotype is on-target. That said, failure to rescue is not always proof that the phenotype is off-target since endogenous expression levels cannot reliably be matched by reexpression from a plasmid which can lead to either too little or too much protein being reexpressed. Depending on the protein, incorrect levels (high or low) can in themselves present with a phenotype.

In order to investigate the role of the RAB7 effector RILP in dendritic degradative pathways, we obtained two previously used sh-plasmids. In our hands, shRILP#1 gives strong phenotypes in cultured hippocampal neurons, including disruption of endosomal compartments and of degradative pathways (Table 1). Unexpectedly, this same sh-plasmid also leads to profound disruption of Golgi and TGN markers, a phenotype not previously described in other cell types. We tested a second shRILP (shRILP#2) which gave none of the endosomal phenotypes observed for shRILP#1 but shared the LAMP1 and Golgi/TGN phenotypes (Table 1). Could this then be a novel on-target phenotype for loss of RILP? For two reasons, we suspect that the Golgi phenotype observed in neurons for shRILP#1 is off-target. 1) We were not able to rescue the shared Golgi phenotypes of shRILP#1 and #2 by re-expression of RILP. Re-expressing RILP to near endogenous levels would be required to cleanly interpret the lack of rescue, something that is not achievable in a transient expression experiment of primary neurons. We note that overexpression of RILP per se did not affect intensity of GM130 staining in neurons. 2) We did not observe any Golgi phenotypes with two other approaches to downregulate RILP, siRILP or Cas9/gRILP. It is thus likely that these phenotypes are off-target. Since the disruption of the Golgi might lead secondarily to disruption of endosomal trafficking, it is not possible to decisively conclude that the observed endosomal phenotypes are on-target for RILP downregulation in neurons. Curiously, the disruption of GM130 and TGN38 did not lead to reduced levels of membrane receptors, such as NSG2. It is possible that the reduced LAMP1 staining is in fact due to disrupted Golgi function. Additional approaches are needed to evaluate the true loss-of function-phenotypes for RILP in neurons.

**Table 1:** Summary of phenotypes reported in this paper.

|  | *neurons* |  | *NRK cells* |  |
| --- | --- | --- | --- | --- |
| *phenotype* | **shRILP#1** | **shRILP#2** | **shRILP#1** | **shRILP#2** |
| **Nsg2 accumulation** | increased | normal | NA | NA |
| **Rab7 staining** | increased | normal | normal | normal |
| **EEA1 staining** | greatly increased | normal | normal | normal |
| **VPS35 staining** | increased | normal | normal | normal |
| **LAMP1 staining** | decreased | decreased | normal | normal |
| **soma size** | normal | decreased | NA | NA |
| **GM130 staining** | greatly decreased | greatly decreased | normal | normal |
| **TGN38 staining** | greatly decreased | greatly decreased | normal | normal |

Secondly, we find that shRILP#1 and shRILP#2 share some suspected off-target phenotypes (Table 1), but not others. Most notable was the strikingly decreased soma size in shRILP#2 but not shRILP#1 expressing neurons. These neurons had the appearance of less mature cells with smaller soma size and shorter dendrites. We also did not observe changes in soma size for siRILP or gRILP/Cas9, suggesting that this is an shRILP#2-specific off-target phenotype. It is not known what the shared and distinct off-target mechanisms are for these two shRILP plasmids.

Since RILP also interacts with RAB34 which is Golgi-associated (12), we tested if dominant-negative RAB34 might phenocopy the loss of Golgi markers. High expression of DN-RAB34 led to fragmented Golgi but not loss of Golgi markers. The levels of the lysosomal protein LAMTOR4 was somewhat reduced as well. We thus discovered that RAB34 plays a role in neuronal Golgi morphology, but if this is via RILP remains to be established. Interestingly, we observed some differences to the reported lysosomal clustering by overexpression of RAB34 in non-neuronal cells: in neurons, lysosomal distribution is not affected.

Lastly, we tested whether Golgi disruption was also observed in non-neuronal cells for shRILPs since such phenotypes had not previously been reported. We found that none of the neuronal phenotypes could be observed in NRK cells (Table 1). Surprisingly then, shRILP plasmids appear to cause specific off-target phenotypes in neurons which are not observed in non-neuronal cells. It is formally possibly that the fact that NRK cells keep dividing and diluting shRILP plasmids whereas neurons do not, might contribute to the observed phenotypic differences. Nevertheless, the fact that the loss of Golgi markers in neurons cannot be rescued and is not phenocopied with siRILP or gRILP-Cas9 leads us to suspect neuronal off-target effects. It is important to point out that with the right controls (such as rescue), short hairpin-mediated downregulation can be used effectively to probe the cellular roles of proteins. Many examples exist in the literature where careful controls were carried out, most importantly rescue experiments (see for example 5, 23–26). Our findings reported here suggest that cellular context can lead to cell-type specific off-target effects. The mechanism of the neuronal specificity of the suspected off-target phenotypes is not known. We recommend that even previously validated knockdown constructs need to be re-validated for use in a different cell type.

## Experimental Procedures

### Materials

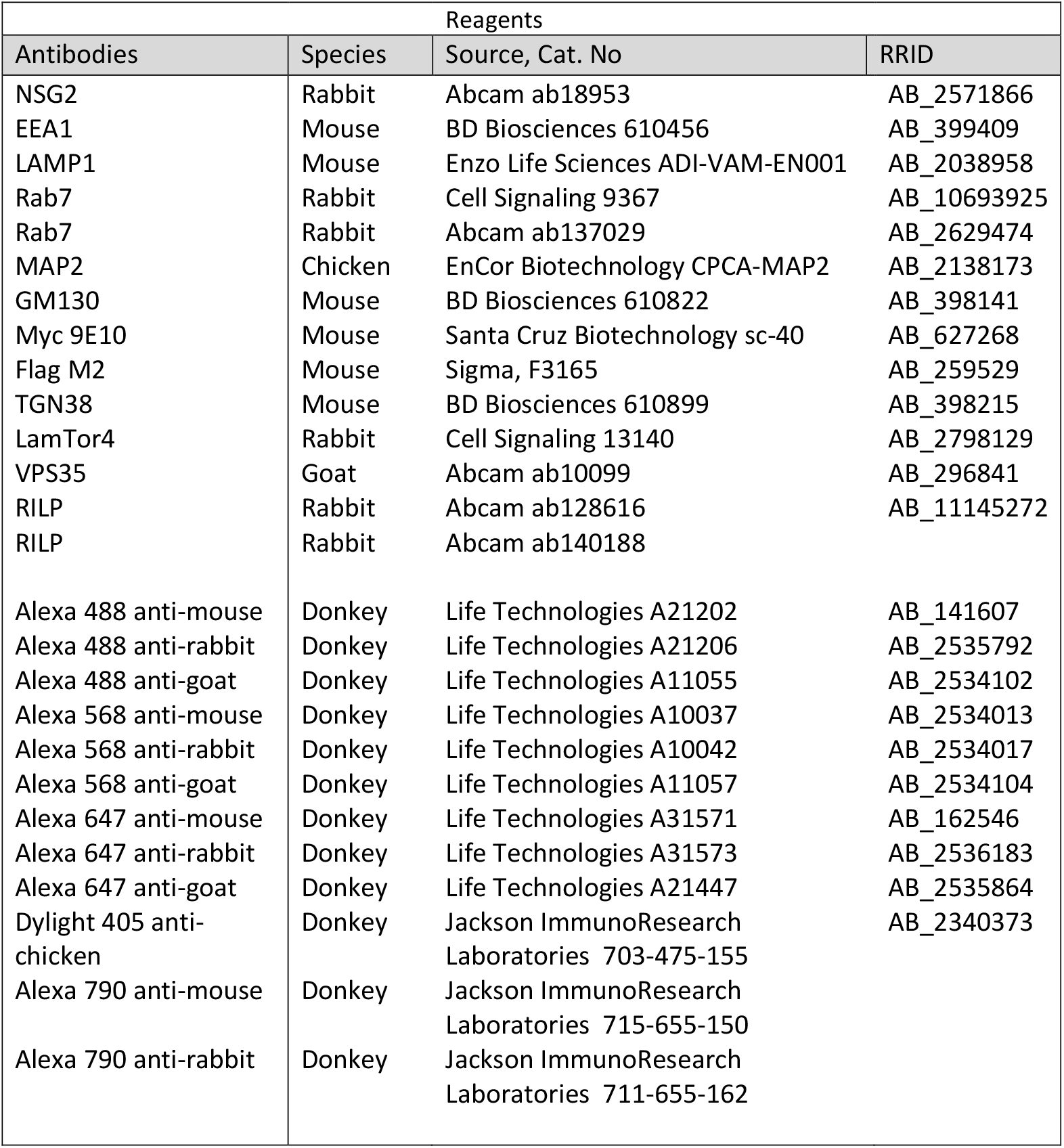
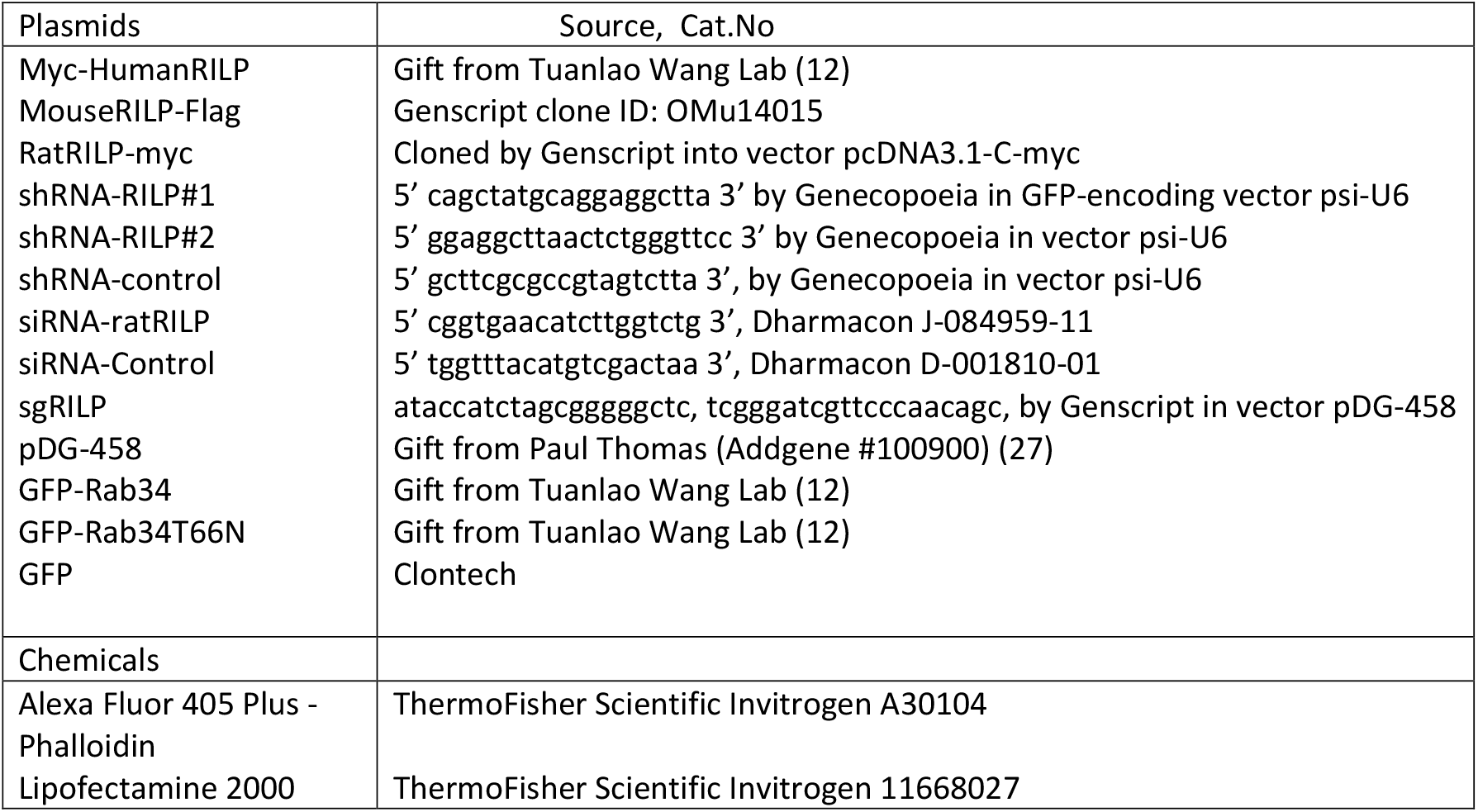

### Neuronal Cultures, mammalian cell lines and transfection

1) Neuronal cultures were prepared as described in (28). In brief, the cultures were prepared from E18 rat hippocampi, as approved by the University of Virginia Animal Care and Use Committee. All experiments were performed in accordance with relevant guidelines and regulations (ACUC protocol #3422). Hippocampi from all pups in one litter were combined and thus contained male and female animals. Cells were plated on poly-L-lysine coated coverslips and incubated with plating medium containing DMEM medium with 10% horse serum. After 4 h, the plating medium was removed and replaced with serum-free medium supplemented with B27 (ThermoFisher), and neurons were cultured for 5–10 DIV (days *in vitro*) for experimental use. Transfections were carried out using Lipofectamine 2000 (Invitrogen). Neurons at DIV7-8 were transfected with either myc-HmRILP, RtRILP-myc, MsRILP-Flag, for 36–40 hours. For RILP knockdown experiments, neurons at DIV5-6 were transfected with shControl, shRILP#1, shRILP#2, Cas9 and GFP gene-containing empty vector pDG458, gRILP, or siRILP mixed with the GFP plasmid for 6 days. For RILP rescue experiments, neurons at DIV5-6 were transfected with shControl, shRILP#1, or shRILP#2 together with myc-HumanRILP for 6 days. To test the knockdown efficacy of gRILP, neurons were transfected at DIV5 with gRILP, or vector pDG-458 along with RatRILP-myc for 6 days. To investigate the effects of RILP knockdown on NSG2 degradation, transfected neurons were incubated with/without cycloheximide (CHX, 20μg/ml) for 4hrs. All transfection experiments were repeated in at least 3 independently derived cultures.

2) HEK293 and NRK cells were maintained in DMEM+10%FBS. The cells were transfected with myc-HumanRILP, MouseRILP-Flag, or RatRILP-myc for 48hours using Lipofectamine 2000. To investigate the effects of RILP knockdown on endosomes/organelles in mammalian cells, NRK cells were transfected with shControl or shRILP#1 for 2 days, followed by trypsinization and replating the cells for another 3 days. All transfection experiments were repeated in at least 2 independently derived cultures.

### Immunocytochemistry

Immunostaining of neurons was carried out as described in (4, 28). Neurons were fixed in 2% paraformaldehyde/4% sucrose/PBS in 50% conditioned medium at room temperature for 30 minutes, quenched in 10 mM glycine/PBS for 10 minutes. After washing with PBS, cells were then blocked in 5% horse serum/1% BSA/PBS ± 0.2% TritonX-100 or 0.1% saponin for 20 minutes. All antibodies were diluted in 1% BSA/PBS and incubated for 1 hour. Coverslips were mounted in Prolong Gold mounting medium and viewed on a Zeiss Z1-Observer with a 40x objective (EC Plan-Neofluar 40x/0.9 Pol WD = 0.41). Apotome structured illumination was used for most images. Images were captured with the Axiocam 503 camera using Zen software (Zeiss) and processed identically in Adobe Photoshop. No non-linear image adjustments were performed.

### Western Blot

HEK293T cells were transfected with human myc-HmRILP, mouse MsRILP-Flag or rat RtRILP-myc for 3 days and harvested for western blot analysis after lysis in lysis buffer (Cell signaling #9803). Blots were imaged with Li-Cor Odyssey CLx Imager. All western blot analyses were repeated twice.

### Quantification of Soma Intensity

Soma intensities were quantified using Imaris 9.5.1. Briefly, the transfected cells were identified, the somas were masked and the average intensity was measured after background correction. Between 20-30 cells per experiment were quantified for 2-3 independent experiments.

## Data Availability

The data described are contained within the manuscript.

## Supporting Information

This article contains supporting information.

## Acknowledgments

We thank Dr. Tuanlao Wang (Xiamen University) for generously providing the Rab34 plasmids. We thank all members of the Winckler Lab for critical feedback on this work.

## Funding and additional information

This work was funded by NIH R01NS083378 (to BW). The content is solely the responsibility of the authors and does not necessarily represent the official views of the National Institutes of Health.

## Abbreviations

RILP: Rab interacting lysosomal protein
CHX: cycloheximide
DIV: days in vitro
EE: early endosome
LE: late endosome
sh: short hairpin
LAMP: lysosome-associated membrane protein
TGN: trans-Golgi network
NRK: normal rat kidney
HEK: human embryonic kidney
DN: dominant-negative
NSG2: neural-specific gene 2

