## Supplement for "“Disruption of Golgi markers by two RILP-directed shRNAs in neurons: a new role for RILP or a neuron-specific off-target phenotype?”"

List of supporting material:

1) Table S1: Comparison of knockdown approaches in the published literature
2) Figure S1: Abcam anti-RILP antibodies ab128616 and ab140188 recognize human, but not rat or mouse RILP.

#### Supplementary Table 1: comparison of RILP knockdown approaches in the published literature

| paper | cells | species | Interference (# of indep. reagents) | sequence | % efficacy (RILP levels) | phenotypes |
|---|---|---|---|---|---|---|
| Progida et al **(ref 13)** | HeLa | human | siRNA (2) | *siRNA-RILP1, sense sequence 5 - GCAGCGGAAGAAGAUCAAGTT-3 and antisense sequence 5 - CUUGAUCUUCUUCCGCUGCTT-3 ; siRNA-RILP2, sense sequence 5 - GAUCAAGGCCAAGAUGUUATT-3 and antisense sequence 5 - UAACAUCUUGGCCUUGAUCTT-3* | >80% (WB) | elevation of 4 LE markers (LAMP1., CD63, LBPA, M6PR)<br><br>EGFR degradation delayed in EE |
| Zhou et al. **(ref 8)** | Min6<br>Mouse insuloma<br><br>Ins-1<br>Rat insuloma | Mouse<br><br><br>rat | shRILP (3)<br><br><br>gRNA/Cas9 (1)<br><br>Cell penetrating peptide (2) | #1: cagctatgcaggaggctta (same sequence as rat. This is our shRILP#1)<br>#2: cagagcttggaacctgatg<br>#3: gtccaaggtgtttctgctg<br><br>*5′-GCCACTAGTAGTGCGGGCGC-3′*<br><br>CPP-RILP-1 (251–262aa: CPP-ILQERRNELKANV),<br>CPP-RILP-2 (297–307aa: CPP-QRRKIKAKMLG) | >80% (WB for #1) | increase pro-insulin levels and increase insulin secretion<br><br>increase pro-insulin levels and increase insulin secretion<br><br>increase pro-insulin levels and increase insulin secretion |
| Ye et al. **(ref 10)** | Superior Cervical Sympathetic neuron cultures | mouse | shRILP (SMART Pool) | CAGCTATGCAGGAGGCTTAAC; this is our shRILP#1<br>AGATCAAGGCCAAGATGTTAG;<br>CCAGAATTTCTTTGGCTTATG;<br>TTCAGCAGGGAAGAGCTTAAG;<br>AGGAGCGGAATGAGCTCAAAG | Not reported. | TrkA signaling endosomes are not transported retrogradely in axon. |
| Khobrakar et al. **(ref 9)** | Cortical neuron cultures. NRK cells. C6 glioma. | rat | shRILP from IDT (1) | 5′-GGAGGCTTAACTCTGGGTTCC- 3′ (this is our shRILP#2) | >60% (WB)<br>~70% (RT-PCR) | decrease in motility of axonal autophagosomes and late endosomes (Rab7)<br>decreased autophagosomal clearance. |
| Cason et al. **(ref 11)** | Hippocampal neuronal cultures | rat | siRILP (1) (ON-TARGETplus; Dharmacon) | 5′-CGG UGAACAUCUUGGUCUG-3 (This is our siRILP) | ~ 25% (WB) (36 hours) | small motility defect of axonal autophagosomes. |

Figure S1: *Abcam anti-RILP antibodies ab128616 and ab140188 recognize human, but not rat or mouse RILP.*
(A-F) DIV9 rat hippocampal neuronal cultures were stained with anti-RILP antibodies ab128616 (A,C,E) or ab140188 (B,D,F) and anti-myc antibody (A-D) or anti-FLAG antibody (E,F) after 48 hours of expressing human myc-RILP (A,B), rat RILP-myc (C,D) or mouse RILP-FLAG (E,F). Both anti-RILP antibody detected human myc-RILP but not rat RILP-myc or mouse RILP-FLAG.
(G,H) HEK293 cells were transfected with human myc-RILP (lane 1), mouse RILP-FLAG (lane 3) or rat RILP-myc (lane 4). Untransfected HEK203 lysates were loaded in lane 2 as controls. Lysates were probed with ab128616 (G) or ab140188 (H). The arrow points at human myc-RILP. No band is recognized at the expected size for mouse or rat tagged RILP. The correct sizes of human myc-RILP, rat RILP-myc and mouse RILP-FLAG can be seen on the blots in (I).

### Supplementary Figure 1

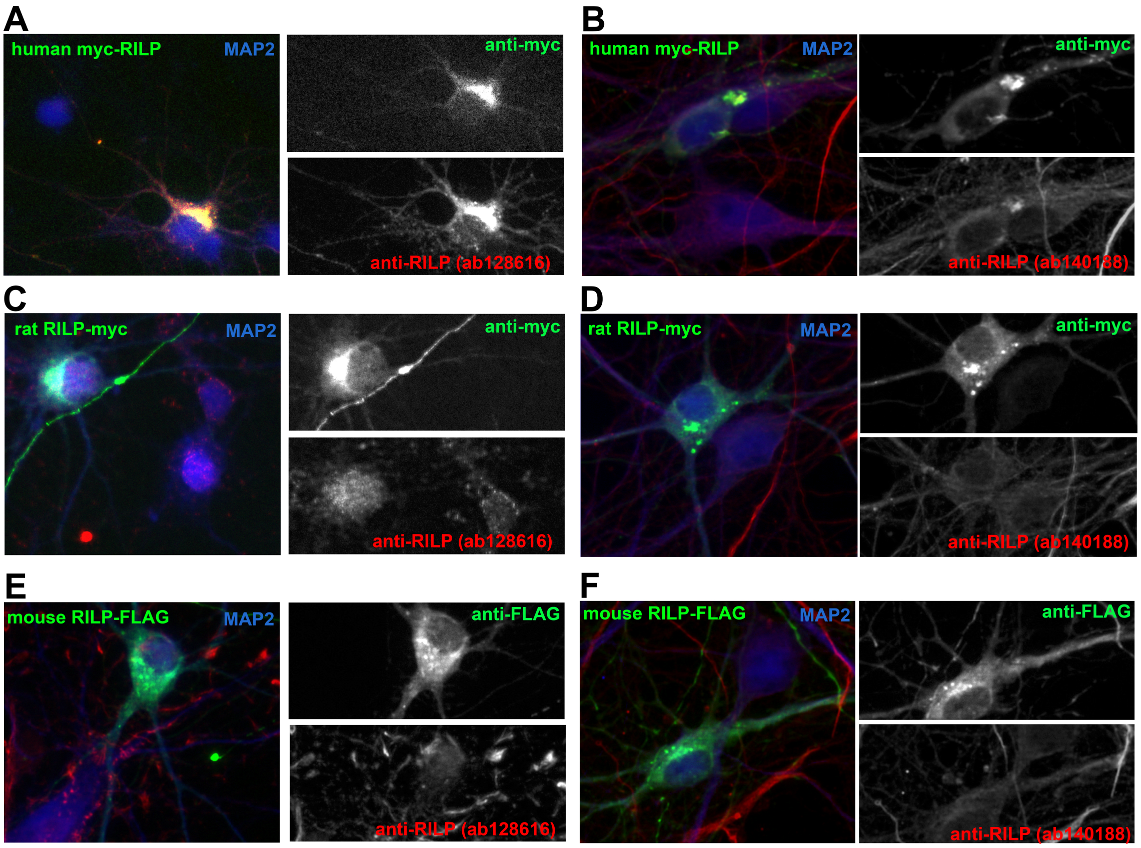


#### Western blots on HEK293 lysates

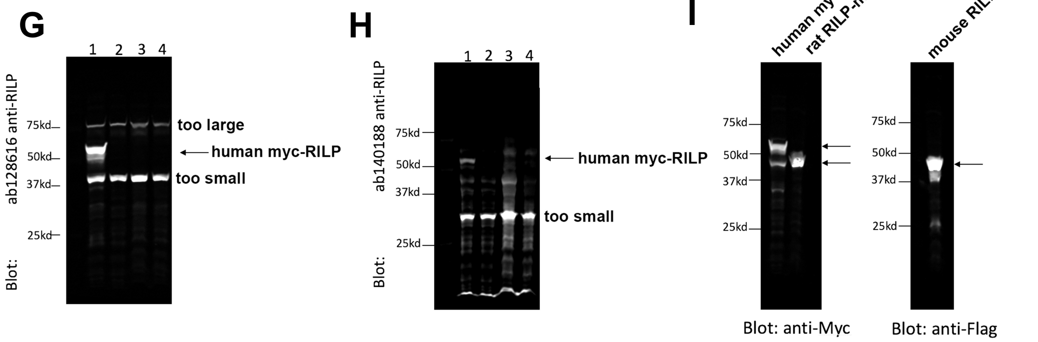


1: 293 cells transfected with myc-humanRILP
2: non-transfected 293 cells
3: 293 cells transfected with mouseRILP-Flag
4: 293 cells transfected with ratRILP-myc
